# *EagleImp*-*Web*: A Fast and Secure Genotype Phasing and Imputation Web Service using Field-Programmable Gate Arrays

**DOI:** 10.1101/2022.02.24.481790

**Authors:** Lars Wienbrandt, Christoph Prieß, Jan Christian Kässens, Andre Franke, Franziska Uhing, David Ellinghaus

## Abstract

Imputation web servers have been developed that allow phasing and imputation of genome-wide data without the need for own computing resources. However, their terms of use, data sharing with third parties, the security architecture of the web service, and the algorithms and parameter settings used are only partially disclosed. We developed *EagleImp-Web*, a fast, secure and convenient web service for phasing and imputation of genome-wide data. The web service uses technical improvements in phasing and imputation algorithms and a *field-programmable gate array (FPGA)* accelerator design to reduce computation time without loss of phasing and imputation quality. Other key features include no exposure of user information and input/output data to third parties, high data security and fast secure download, user authentication through 2-factor authentication, full control in managing user accounts, and full transparency of algorithms and their settings. *EagleImp-Web* provides simple and convenient functionalities for monitoring running jobs and selecting parameter settings and output information. Due to the speed advantage over a purely CPU-based implementation, *EagleImp-Web* offers the user the ability to choose a more resource-intensive parameter setting in exchange for computation time to further improve phasing and imputation quality. *EagleImp-Web* is freely availabe at https://hybridcomputing.ikmb.uni-kiel.de.

## Introduction

Imputation web servers have been developed that allow researchers without own computing resources to easily perform genome-wide phasing and imputation with reference panels from different genome sequencing projects [1]. New whole genome sequencing (WGS) projects will lead to huge reference panels in the near future, such as that of the European 1+ Million Genomes Initiative (1+MG). To keep phasing and imputation practical and runtimes short, we recently optimized current phasing (*Eagle2* [2]) and imputation software (*PBWT* [3]) and combined both steps into a single freely available application named *EagleImp*, resulting in a speed advantage of two to ten-fold over the original tools with improved phasing and imputation quality [4].

Here we introduce *EagleImp-Web*, a fast and convenient genotype phasing and imputation web service that provides all the features of *EagleImp* free of charge to the community. *EagleImp-Web* complies, due to its security and data protection architecture, with the European Union (EU) General Data Protection Regulation (GDPR). The terms of the GDPR are explained transparently on our website. The service runs on university computers located in Kiel, Germany. All website functions are available via an encrypted and certified *https* connection. The website does not store cookies besides necessary session cookies on the user’s computer and is free of advertising and does not collect any data for marketing purposes or for sharing with third parties. For job submission and management functions, a user login with only minimal requirements is required, i.e. a valid email address and password. The email address is required for notification messages about the user’s jobs and is also used for password recovery. File downloads are handled directly on our server via certified *https* and can be started either directly from the protected user account or with a one-time password from the command line (e.g. via the download tool *wget*). This eliminates the time consuming encryption and decryption of the result files. Compared to other imputation web servers, input and output data is stored temporarily and exclusively on our university computers in Kiel, Germany, and are never passed on to third-party services (such as *Globus* or *Amazon* servers, or some unknown location). Data access is only possible for the authorised user. Additional security is provided by optional 2-factor authentication with the option to register different devices (e.g. a smartphone with fingerprint authentication or face recognition, or a USB security key).

To further reduce computation times through the web service, we have also re-implemented the phasing core algorithm of *EagleImp* [4] in *EagleImp-Web* by using a field-programmable gate array (FPGA), which offers a further speed increase of up to 66% with no loss in accuracy when compared to the CPU-only approach of *EagleImp*. The FPGA accelerator is seamlessly integrated into the computing system of our web service and utilised transparently for the user. The speed advantage allows, for example, the use of a higher-value parameter selection for phasing with an improvement in the phasing quality while maintaining the same runtime.

## Materials and Methods

The server infrastructure of *EagleImp-Web* is divided into two components, referred to as *frontend* and *backend* system, to make our service fast, stable and secure (**Figure 1**). It is described in the first part of this section, while in the second part we explain the basic concepts of the underlying *EagleImp* software and the integration of the FPGA accelerator.

**Figure 1.**
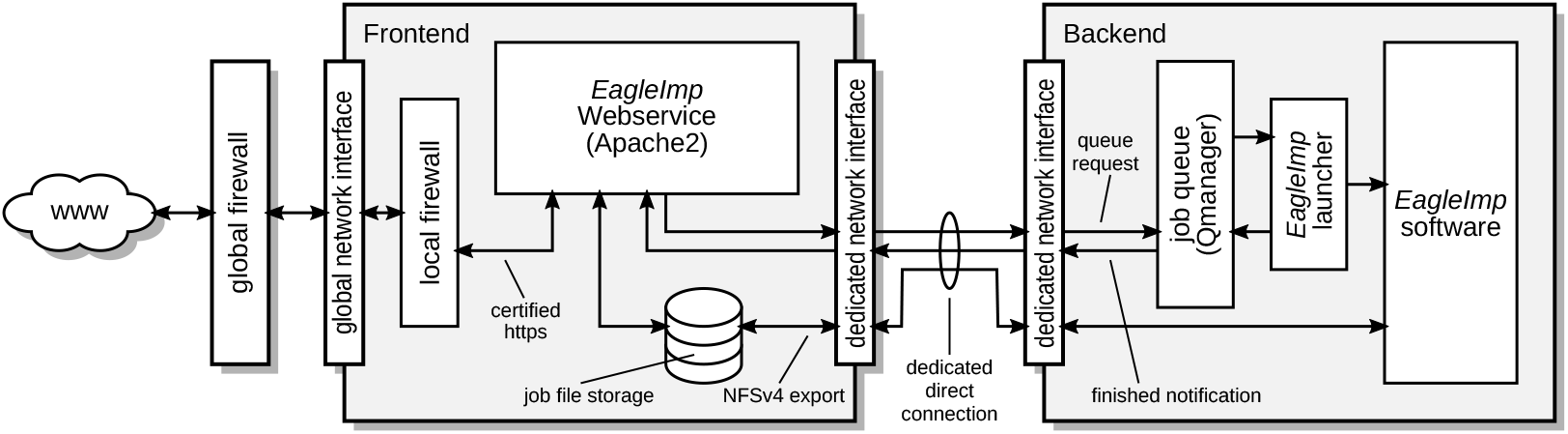
The *EagleImp-Web* server infrastructure consists of two components: frontend and backend system. The web service is hosted by the frontend while jobs are processed on the backend system. The processing order is controlled by the job queue which we implemented in the *Qmanager* software on the backend. Files are stored on the frontend and accessed via *NFSv4*. The division into frontend and backend system allows job processing to be independent from web service operations and keeps the web service functional despite a full computational load on the backend during job processing.

### *EagleImp-Web* Server Infrastructure and Functionality

The *frontend* hosts the web service including a database with user information, such as login credentials, submitted jobs and their results. Uploaded files and result files are also stored on this system. The frontend is equipped with an Intel Xeon Silver 4110 8-core CPU @ 3 GHz and 128 GB RAM. It offers more than 7 TB of redundant storage capacity and is currently running on an Ubuntu 21.10 Linux system, which is regularly updated. The web service is implemented in *PHP* and *JavaScript* and is hosted by an *Apache2* server (currently version 2.4.46). The database is implemented in *PostgreSQL*, version 13.5.

With two Intel Xeon E5-2667 v4 8-core CPUs running at @ 3.6 GHz and 256 GB RAM, the *backend* provides the necessary computing power for the actual processing of the submitted jobs. Ubuntu Linux 21.10 is also used as the operating system. The backend is further equipped with an FPGA accelerator, an Alpha Data ADM-PCIE-8K5 PCIe card containing a Xilinx Kintex UltraScale KU115 FPGA. The backend’s main tasks are to host the job queue, that we have newly implemented in our *Qmanager* [5] software, and to run the *EagleImp* [4] tool to process the user’s phasing and/or imputation jobs.

To ensure direct communication without potential interception risks and routing problems, the frontend and backend are connected via a direct Ethernet connection and dedicated network devices. Communication between these systems via this connection is limited to two applications: First, the backend offers access to the *Qmanager* via a single open port on this direct connection to the frontend. Second, the frontend only exports the storage file system via *NFSv4* to the backend via this connection and accepts notification calls from the *Qmanager*. A firewall on the frontend (implemented with the Linux system tool *iptables*) rejects all incoming traffic from the internet except *https* requests. (*http* requests are also allowed, but are automatically redirected to *https*). The backend firewall is configured to completely block all incoming traffic from the internet. Only administrative access via *SSH* to the local network is exceptionally allowed for both systems. Since the servers are located in the infrastructure of Kiel University, an additional firewall from the university’s router ensures that rules are not violated. The server’s certificate (required for secure *https* connections) is issued by the external organization *Deutsches Forschungsnetzwerk* (DFN; German Research Network) and will be verified by any browsers standard certificate chain.

Since the frontend makes the file system available to the backend via *NFSv4*, the web service can directly access all outputs of the *EagleImp* software running on the backend. The *EagleImp* software continuously updates its progress in the status files observed by the web service, so that the web service user can constantly monitor her/his running job. Global job management is done through our newly developed job queuing system, which is implemented in the *Qmanager* software hosted on the backend system [5]. Its main function is to receive commands to be processed in the backend system. It processes all queued, running and finished jobs (including their output and return codes) from the frontend system. A finished job is reported directly to the web service via a notification script that triggers several operations on the frontend system, such as changing the job status, notifying the user and preparing the download URLs for the results. For more details on the implementation of *EagleImp-Web*, we refer to **Supplementary 1**.

### *EagleImp* Software and FPGA Implementation

The *EagleImp* software is freely available at *GitHub* [6]. It is based on the popular phasing tool *Eagle2* [2] and imputation tool *PBWT* [3], and combines both steps in a single application. The main advantages of *EagleImp* over the classical two step approach with *Eagle2* and *PBWT* are the increased computation speed of a factor 2 to 10 while the phasing and imputation quality is at least maintained or even improved. The speedup results from introducing multiprocessing with several temporary output files for imputation reducing the input/output (IO) bottleneck among other algorithmic improvements regarding interval mapping in the internal *Position-based Burrows-Wheeler Transform (PBWT)* data structure and the extensive usage of bit operations and processor directives. For more details on the basic concept of phasing and imputation in *EagleImp*, we refer to Wienbrandt *et*.*al*. [4], also summarized in **Supplementary 2**.

To further accelerate the web service, we introduce support for an accelerated phasing step in *EagleImp* here using *field-programmable gate arrays (FPGAs)* [7, 8]. An FPGA is a processing unit where the logic circuits in the hardware are programmable which is completely contrary to *Central Processing Units (CPU)* which are commonly known as “processors”. The main advantage of FPGA computing is that simple operations, especially those based on Boolean logic, can be implemented directly in the circuit’s logic which makes them extremely fast and keep the power consumption to a minimum. Furthermore, by implementing many operations in a pipeline chain or in parallel, many operations can be performed concurrently. The developer can flexibly design the FPGA core with arbitrary word sizes and parallel data channels to implement a core algorithm with the goal of an extremely optimized use of the devices resources. In contrast, CPUs lack this flexibility as they process a “program” which actually is a chain of subsequent commands. Parallelism is introduced only by multiple CPU cores and pipelining for certain commands, but word sizes and data channels are fixed.

In *EagleImp* many operations in the phasing preliminaries (such as selecting the *K*-best haplotypes and building the condensed reference as well as the creation of the PBWT) are based on Boolean operations. Therefore, we outsourced this part into an FPGA design, illustrated in **Figure 2**. In contrast to our first approach presented in [9], we have improved the design by adding an entity that selects the *K*-best haplotypes for each target from the reference (*K* is a user option and defaults to *K* = 10, 000 in *Eagle2*), and we have implemented the generation of the PBWT from the condensed reference on the FPGA (forward and backward PBWT as required for the forward and reverse phasing steps). We were able to remove the matrix transposition step from our previous FPGA design as the entire pipeline can now be processed in variant-major format without an intermediate conversion to a sample-major format. (The variant-major format is also used in VCF files.) For details on how to integrate the new FPGA design into the phasing process, see **Supplementary 3**. Note that FPGA acceleration is only possible for *K ≤* 32, 768 due to the limited *block RAM (BRAM)* resources on the FPGA. Runs with a higher *K* parameter are always performed without FPGA acceleration.

**Figure 2.**
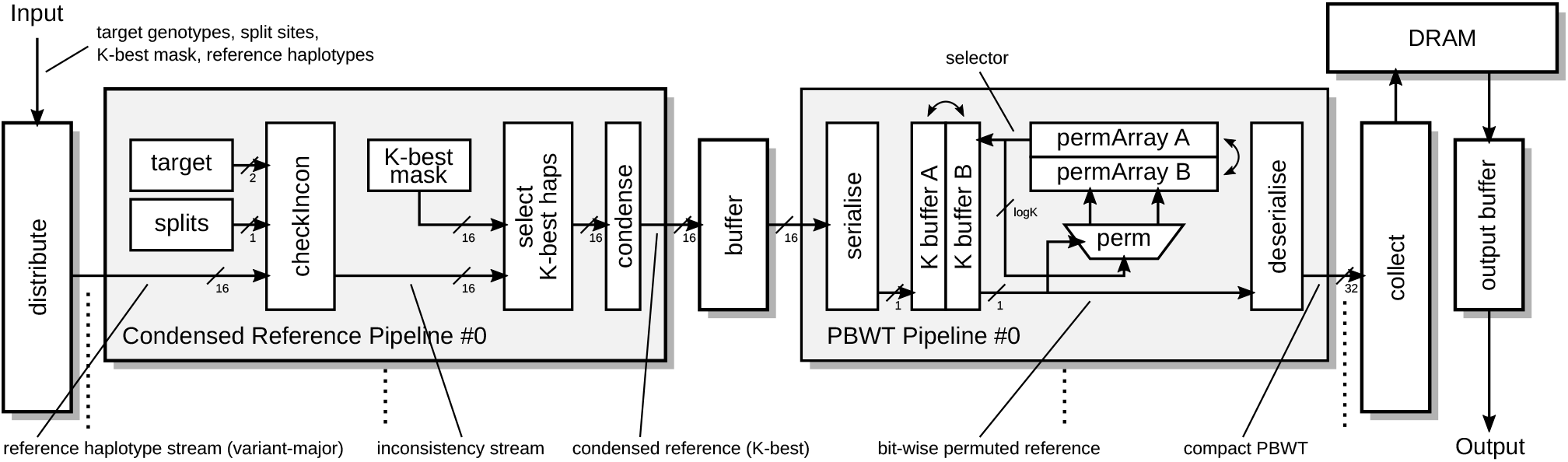
The FPGA pipeline design to create the condensed reference and the PBWT data structure for the phasing step. For simplicity, the figure shows 1-bit processing per clock cycle. In contrast, the implemented design processes 2 bits in one clock cycle using dual-port BRAMs and doubling data, address, permutation, multiplexer and output routes. For the basic concept of phasing using FPGAs, we refer to Wienbrandt *et*.*al*. [9]. Details of the current FPGA implementation and an explanation of the illustrated design are provided in **Supplementary 3**.

## Results

### User Interface and Functionality

#### Registration and Login

Use of *EagleImp-Web* first requires the registration of a personal user account, which can be created by providing a valid email address and password (optional 2-factor authentication with different devices is available for additional security, see below). The email address is used to verify the account, as a confirmation email is sent to the user upon registration. The user account enables protection against unauthorized access by third-parties to imputation results and a variety of convenience features in connection with our service. We explicitly point out that we do not use the provided email address for purposes other than job notifications and account management and do not collect any usage information or statistics of our service in connection with user accounts. After login, it is possible to submit jobs, manage ongoing or completed jobs, download results, and manage accounts. The web service will automatically logout the user after 30 minutes of inactivity.

#### Job Submission

New jobs are arranged in a queue to ensure a fair order of execution among users (on a first-come, first-served basis). To run a job within our service, the upload of target genotype data (to be phased and/or imputed) in *Variant Call Format (VCF)* is required. Accepted file formats are .vcf.gz and .bcf, and uploaded files are subject to the following restrictions: Only one file per chromosome may be uploaded, the filename must start with the chromosome number with an optional leading “chr”, the samples must be the same in all uploaded files, and the sample limit is 100,000. The consistency of the files is checked during upload.

Figure 3. shows a screenshot of the job submission form. Uploading the target files is possible in three different ways. The easiest way is to upload via a browser. Files can simply be selected in a file selection dialog or dragged and dropped in the designated box. After clicking on “Submit Job”, the upload is starting. Alternatively, the web server can actively fetch the files from public URLs (e.g. pointing to a private server) that are provided in the submission form. Or, upload via *Secure File Transfer Protocol (SFTP)* can be chosen, which requires the user to provide the host URL, login credentials and relative paths to the files on that server. Note that we use the login credentials only for the purpose of downloading the files. We never submit them in plain-text, as SFTP is an encrypted connection, and we delete them immediately after the download to our server has finished.

**Figure 3.**
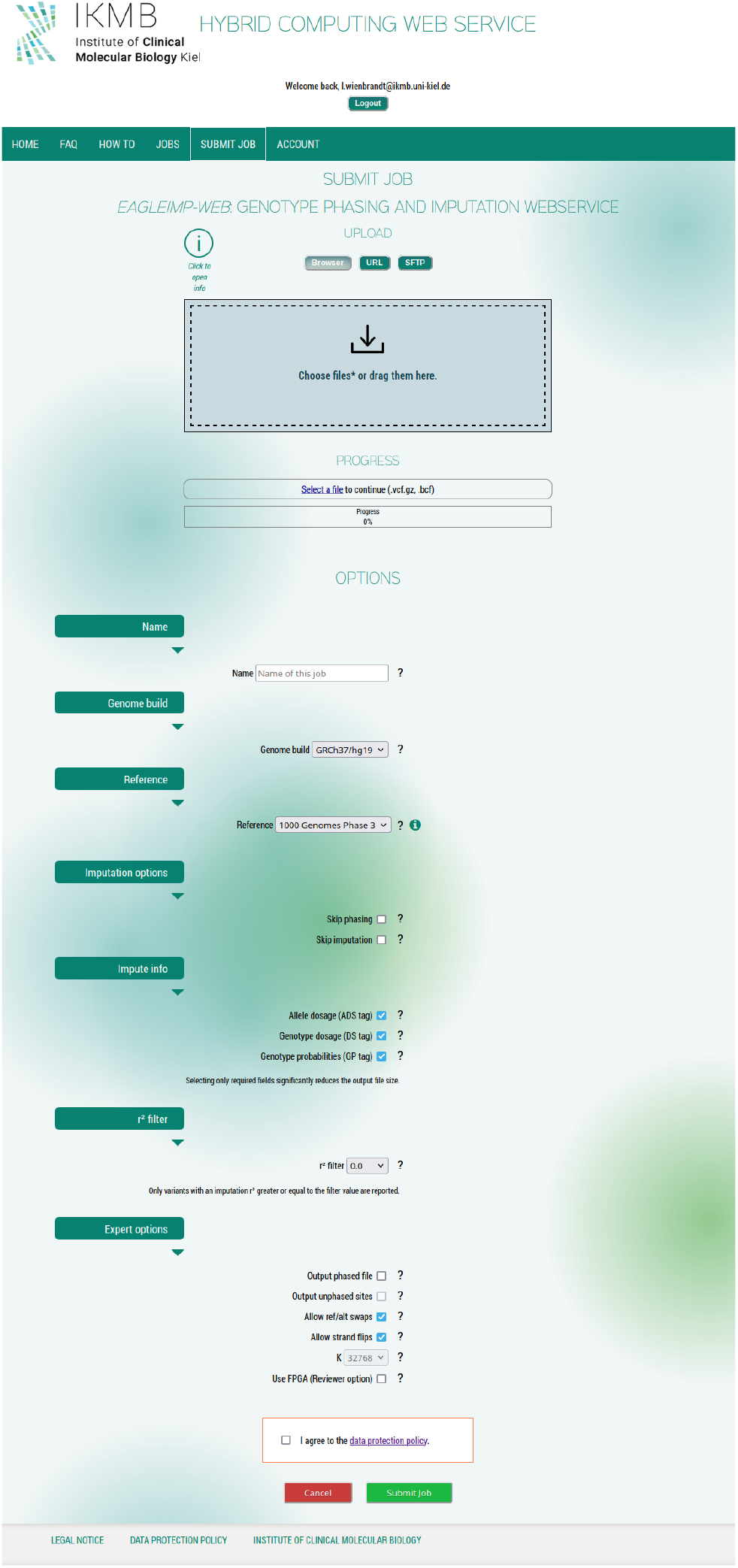
Screenshot of *EagleImp-Web*’s job submission page with the available configuration options.

The runtime options of the job include an arbitrary job name, the genome build of the input data (either *GRCh37* or *GRCh38*) and the reference panel (currently either the *1000 Genomes Phase 3* panel [10] or the *European Genome-Phenome Archive (EGA)* release of the HRC1.1 reference panel [1, 11] are available — as both panels were released in GRCh37, we provide lifted versions of both panels for GRCh38 imputation (see **Supplementary Table 1** for more details on the available reference panels).) If the user has prepared a pre-phased dataset for upload, the phasing process can be skipped. Likewise, the user can skip the imputation process if a phasing-only run is desired. Other parameters include the option to select the type of dosage information that will be reported along with the imputed genotypes (hard calls), i.e. the allele dosages, the genotype dosage, the genotype probabilities, a combination of those three or none. The user can select an imputation accuracy *r*^2^-filter threshold to reduce the imputation output to variants of a minimum quality. Other expert options include handling *ref/alt swaps* or *strand flips*, and the *K* parameter can be chosen from three different categories (only available if the reference panels contains more haplotypes than the smallest *K*-parameter).

#### Job Management

After submission, the job is queued in the job queue and can be managed in the “Jobs” section. The job management page is divided into four parts: queued jobs, running jobs, finished jobs and deleted jobs. The jobs are listed in chronological order along with a waiting status indicating when the job will be processed. A single user can queue up to three jobs at a time (i.e. no more than three jobs can be queued waiting to be processed by a single user, however, there is no limit to the total number of jobs processed for a user).

Figure 4. shows a screenshot from an exemplary job progress. Once a job is actively being processed, the user can monitor the current progress and view information about each file, including warning and error messages. When a job is regularly completed, it is moved to the finished jobs section from where the result files can be downloaded. A job can also be canceled, in which case it is stopped and also moved to the finished jobs section.

**Figure 4.**
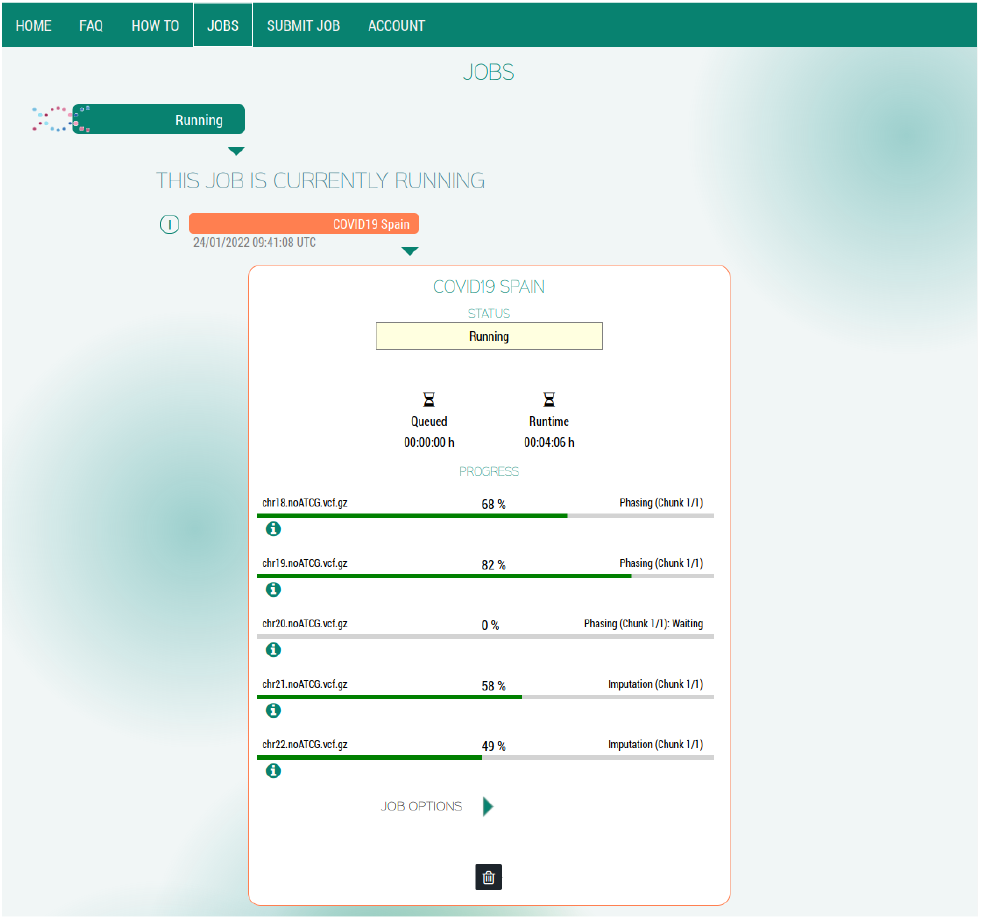
Screenshot of an exemplary imputation job progress in *EagleImp-Web*.

#### Results Download

For each uploaded input file the web service generates the following output files: Imputation results are found in .imputed.vcf.gz files and contain all imputed variants (phased genotypes; hard calls) of the selected imputation reference and optionally the user selected information on allele dosages, genotype dosages and/or genotype probabilities. For each variant, the user receives additional information such as imputation accuracy (*r*^2^), minor allele frequency (*MAF*), reference panel allele frequency (*RefPanelAF*), allele count (*AC*) and allele number (*AN*). Variant IDs and allele coding are taken from the reference panel, and variants that were already present before imputation are marked as TYPED. If the user selects to save the phasing output separately, the web service generates a .phased.vcf.gz file containing the phased input variants. A .phased.confidences file contains the average phasing confidence for each input sample. In addition, the *EagleImp* log file .log is provided, and an info file .varinfo contains separate information for each input variant about how it was included in the analysis (e.g. with a ref/alt swap) or, if it was not included, why it was excluded from analysis (e.g. if it was multi-allelic).

The result files are registered in our database with a unique random string for each file. The web service then generates a secure download link for each file over an encrypted *https*-connection based on this random string and an individual script to download all available files at once. For all completed, cancelled or otherwise terminated jobs, the user receives a notification email containing a generated password uniquely associated with that job to download the results via the command line. The files are locked by default and a direct download via the file URL is only possible when the user is either logged-in to download individual files directly from the website or the password received in the notification email is entered via the download script on the command line. This password is used to unlock the individual files, thus allowing the download only to those users who have access to the download links with the correct URL and the password. In this way, separate encryption of the result files is not required and saves the user time by not having to decrypt the files before using.

In contrast to other prominent imputation web services, all input files are processed independently of each other and thus the failure of individual input files does not lead to the complete abortion of the job. Result files generated before the termination of the job (regardless of whether it was aborted or successfully completed) will be made available for download so that the user can specifically fix the incorrect files without having to run the job for correct files again. Note that we automatically delete all data related to a job (with the exception of status and log files) from our server 7 days after the job is finished unless the job is manually deleted by the user. We keep status and log files together with the job metadata for the user in the deleted jobs section so the user has a history of her/his previous jobs. Of course, jobs in the job history can also be deleted manually (either individual jobs or all deleted jobs at once). Jobs older than one year are automatically removed completely from the history.

#### Account Management

As stated above, a valid email address and a password are required to set up a personal user account. The password is not stored in plain text, but as a secure SHA-256 hash on our server. *EagleImp-Web* also provides a password recovery function in case the user has forgotten the login password. Optionally, the user can activate 2-factor authentication for her/his account. This can be done by adding at least one security key, e.g. from a USB key dongle, a fingerprint reader (e.g. on a smartphone) or another suitable device. Note that once 2-factor authentication has been activated with the registration of a security key, it is no longer possible to log in without the key. The account settings section further contains basic account management functions, such as changing the email address or password and adding or removing a security key for 2-factor authentication. The user may also delete her/his account in the account settings, which results in an immediate deletion of all data and jobs related to the user from our server without exception.

### Quality and Runtime Performance

In our *EagleImp* software publication [4] we showed that *EagleImp* delivers at least the same or better phasing and imputation quality (in terms of phasing switch error rate and imputation genotype error rate) compared to the tools *Eagle2* [2] and *PBWT* [3], with a speed advantage of a factor between 2 and 10. For common variants examined in typical GWAS studies, we were also able to show that *EagleImp* had the same or higher imputation accuracy (in terms of imputation accuracy *r*^2^) than the *Sanger Imputation Service* [12], the *Michigan Imputation Server* [13] and the most recent *TOPMed Imputation Server* [14], despite larger (not publicly available) reference panels. For more details, we refer the reader to Wienbrandt *et*.*al*. [4].

For *EagleImp-Web* we have developed an FPGA design to accelerate the phasing step in *EagleImp* that returns the same results as *EagleImp*’s CPU-only code. First, we conducted quality assurance benchmarks depending on two different values of the phasing parameter *K* (to select the K-best haplotypes from the reference for phasing) by repeating the benchmarks in [4] for the datasets *HRC*.*EUR, COVID*.*Italy* and *COVID*.*Spain* (see **Supplementary Table 2** for more details on the benchmark datasets) with FPGA acceleration enabled. We confirmed for the *HRC*.*EUR* dataset (comprising 494 European samples) the same phasing switch error rate of 0.00470 for *K* = 10, 000 and 0.00434 for *K* = 32, 768 when phased against the EGA release of the *HRC1*.*1* reference panel. The imputation genotype error rates were confirmed with 0.00265 for *K* = 10, 000 and 0.00254 for *K* = 32, 768, respectively. Consequently, the imputation accuracy *r*^2^ shows the same behaviour as demonstrated by determining the *r*^2^ values stratified by minor allele frequency (MAF) for *K* = 10, 000 and *K* = 32, 768 for the real-world datasets *COVID*.*Italy* and *COVID*.*Spain* (from [15], comprising 2,113 Italian and 1,792 Spanish samples respectively) with the resulting diagrams showing no difference for the respective *K* (**Supplementary Figures 1 (a–d)**).

To determine the speed advantage of the FPGA implementation over the CPU-only implementation, we performed a runtime analysis of the same three benchmark datasets and same *K* values from our quality assurance benchmarks above. Note that the backend system of our presented web service here is the same system as used for the benchmarks in Wienbrandt *et*.*al*. [4], which makes runtimes directly comparable. For FPGA acceleration, we again used the fastest multiprocessor configuration for *EagleImp* reported in [4], i.e. eight worker processes with eight threads each are started simultaneously, with two workers each sharing a lock for multiple exclusion on CPU resources (hereafter referred to as *2×4×8* configuration). Further note that the FPGA is used as a single shared resource for all worker processes via multiple exclusion as well. The CPU-only runs were also performed in *2×4×8* configuration.

For the *HRC*.*EUR* dataset and *K* = 10, 000, the FPGA-run took 18 minutes and 24 seconds, and for *K* = 32, 768, the runtime was 34 minutes and 21 seconds (**Supplementary Figure 2, Supplementary Table 3**). This is an acceleration between 18% and 42% when compared to the CPU-only runs. The (larger) real-world GWAS datasets *COVID*.*Italy* and *COVID*.*Spain* showed an even greater speed-up using the FPGA design, with speedup factors ranging between 1.23 and 1.66 for FPGA-accelerated *EagleImp-Web* when compared to *EagleImp* (**Figure 5, Supplementary Tables 4–5**), which is a speed increase between 23% and 66% when compared to the CPU-only runs.

**Figure 5.**
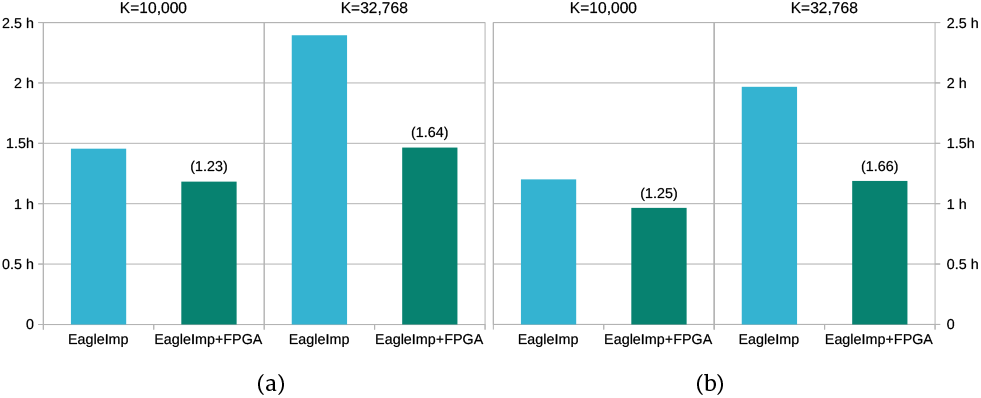
Wall-clock runtimes for two GWAS benchmark datasets (a) *COVID*.*Italy* (2,113 Italian samples with 559,519 variants) and (b) *COVID*.*Spain* (1,792 Spanish samples with 549,696 variants) imputed with *EagleImp* against the HRC1.1 EGA release, using two different values of the phasing parameter *K*. Numbers in brackets indicate the speed increase of *EagleImp* with enabled FPGA acceleration (available in *EagleImp-Web*) versus *EagleImp* without FPGA support.

Because the actual goal of genotype imputation is subsequent genome-wide testing for genetic associations, we examined for the benchmark datasets *COVID*.*Italy* and *COVID*.*Spain* how increasing the *K* parameter affects the genome-wide significant association testing results. In particular, we would like to know, whether the choice of a higher *K* value has a positive effect on the results at the level of association testing. We used *EagleImp-Web* to reproduce our association results from the published GWAS in [15] with three different *K* parameters: *K* = 10, 000, *K* = 32, 768 and *K* = max (where the latter represents the selection of all available reference haplotypes for phasing and is performed with a CPU-only run due to the current FPGA limitation). Phasing and imputation with *EagleImp-Web* using the EGA release of the HRC1.1 reference panel followed by association testing with the *BIGwas* association software pipeline [16] yielded the same two genome-wide significant association signals (at 3p21.31 and 9q34.2) as in [15], whereby for 3p21.31 the *P*-values of association lead variants yielded smaller (i.e. more significant) values when choosing a larger value of *K* (*P* = 5.72 *×* 10–^9^ for *K* = 10, 000, *P* = 1.71 *×* 10–^9^ for *K* = 32, 768, *P* = 9.91 *×* 10–^10^ for *K* = max; **Supplementary Figures 3 (a–c)**). Thus, we assume that improved phasing with the choice of a higher *K* parameter generally also leads to better results in association testing.

## Discussion

Our imputation web service *EagleImp-Web* provides a fast, secure and high-quality service for genome-wide genotype phasing and imputation, with manysecurityand convenience features that other services lack. It is easy to use, freely available, and complies with the European Union (EU) *General Data Protection Regulation (GDPR)*; our GDPR-compliant data protection policy with explicit rights and obligations can be found directly on the website. The web service uses the freely available *EagleImp* software [4, 6], which is based on the popular tools *Eagle2* [2] and *PBWT* [3], but is two to ten times faster with the same or better quality.

In combination with an FPGA accelerator, *EagleImp-Web* is up to 66% faster than using the CPU-only version. Our FPGA-based phasing shows no loss of quality compared to the CPU-only version, but, due to the speed advantage, allows the choice of a higher value for the *K* parameter for phasing, which further reduces the phasing switch error rate, the genotype imputation error rate and increases the imputation *r*^2^, and provides higher quality association results in subsequent genome-wide association analyses (GWAS). Therefore, *EagleImp-Web* offers the user option to select a higher *K*-parameter (*K* = 10, 000 and *K* = 32, 768 with FPGA support, and up to *K* = max without FPGA support — we plan to implement FPGA support for higher values of *K* in the future) to enhance phasing and imputation quality with slightly increased processing time. As an additional feature, users can optionally set expert options to tailor input and output data to their needs, including tolerating or not tolerating Ref/Alt allele swaps or strand flips in the input data, optionally generating a separate file for phasing output alone, and allowing users to select which imputation information (allele dosages, genotype dosages, genotype probabilities) to include in the imputation output.

In contrast to other available imputation web services, *EagleImp-Web* does not require third-party services for file transfers (such as *Globus* in the case of the *SIS*) or unpacking and decrypting files after download (as in the case of the *MIS* and the *TOPMed* services). Files are transferred to and from *EagleImp-Web* via secure connections, a download requires prior user authentication, and download links are unique and known only to the user as exact file locations on the server are replaced by random strings (uniquely associated to a file in the web server’s database). User accounts are password protected and security can optionally be enhanced by 2-factor authentication. For a convenient download of all result files at once, a unique and for each job randomly generated password is used for authentication via the command line.

In addition, job progress and results are displayed transparently to the user, with (e.g. for publication purposes) all selected parameter settings and filter options as well as the exact parameter settings of the underlying phasing and imputation algorithms being reported. Warnings are displayed in case of problems with the input files (e.g. if there are too few genetic variants). Status and progress information is continuously displayed for jobs in progress. If a single file fails, the entire job is not automatically aborted, but an attempt is made to process all remaining input files, and the results and log files are made available to the user for debugging purposes. Our 2-component setup of our service with frontend and backend computer ensures that job processing does not interfere with the web service components. Should we experience a heavy load on our service due to very frequent usage, our setup can be easily scaled with additional backend systems and separate job queues.

Currently, our web service provides the *1000 Genomes Phase 3* reference panel and the EGA release of the *HRC1*.*1* reference panel for phasing and imputation, both as genome build *GRCh37/hg19* and a lifted version to *GRCh38/hg38*, which is different from other web services that perform a liftover of the input data at runtime before phasing and imputation. As a result, the imputation output from *EagleImp-Web* is always in the same genome build as the provided input data. A disadvantage of our first *EagleImp-Web* release is that some reference panels are restricted for general use and have been assembled from dozens of individual projects (such as the *TOPMed r1* panel) or have not been made fully available to the scientific community (e.g. the *HRC1*.*1* reference panel [1]), so we cannot currently provide them as additional panels. However, we are currently working on offering additional reference panels from future genome research projects, especially for previously underrepresented populations.

## Data Availability

The *EagleImp-Web* service can be accessed at https://hybridcomputing.ikmb.uni-kiel.de. All quality and runtime measures from our benchmarks are listed in the **Supplementary Material**. The *EagleImp* software is available at https://github.com/ikmb/eagleimp. The *Qmanager* software is available at https://github.com/ikmb/qmanager. The source code of the FPGA design is available at https://github.com/ikmb/eagleimp-fpga. The *1000 Genomes Phase 3* reference panel can be downloaded at ftp://ftp.1000genomes.ebi.ac.uk/vol1/ftp/release/20130502/. Data access to the EGAD00001002729 dataset (https://ega-archive.org/datasets/EGAD00001002729) for the HRC1.1 panel is restricted and was granted under request ID 11699.

## Acknowledgements

The authors would like to thank the COVID-19 GWAS Group for the use of the COVID-19 GWAS data for the benchmark.

