## Supplementary Material for "*EagleImp*-*Web*: A Fast and Secure Genotype Phasing and Imputation Web Service using Field-Programmable Gate Arrays"

February 24, 2022

**4 Supplementary Tables and Figures** **9**

**5 Exemplary COVID-19 GWAS Imputed with *EagleImp-Web*** **11**

### 1 Implementation of *EagleImp-Web*

#### 1.1 General server configuration

*EagleImp-Web* currently runs on a two systems setup. The first system, the *frontend*, hosts the web server based on *Apache 2.4* [1], a *PostgreSQL* [2] database for user account data and associated files, and data storage. The web service is mainly written in *PHP* and some modules in *JavaScript*. Administrators can configure the service via a separate configuration file. Connections to the service are possible only via encrypted *https*. Our server URL has a valid certificate issued by an external authorization organization (*Deutsches Forschungsnetzwerk*). We renounce the usage of cookies.

The frontend communicates via a direct Ethernet connection with the *backend*. This way no interception by a third-party is possible and the connection is independent of a separate routing system. The backend is a high-performance computing system equipped with two 8-core *Intel*

*Xeon* CPUs and an *Alpha Data ADM-PCIE-8K5* FPGA accelerator card. It hosts the *EagleImp* software [3, 4] and a simple but efficient self-developed job queuing system *Qmanager* [5]. Files are exchanged with the frontend via an *NFS v4* file system that is restricted only to the direct connection between frontend and backend. Also, the communication with the queuing system is done via this connection.

#### 1.2 Account management and user login

In order to use our a service, a personal user account is required. The creation requires a valid email address and a password, but no further information. We do not store plain-text passwords, as it was extremely insecure, but secure *SHA-256* password hashes instead in our database.

Account security can be improved with 2-factor-authentication based on the recent standard *Web Authentication API (Webauthn)* [6]. Webauthn enables usage of hardware authenticators with public and private key-based credentials to perform an SSL handshake between server and the client's authenticator as trusted device. A trusted device may be a USB key dongle, fingerprint reader or facial recognition on a phone or any other applicable device. In detail, an account which is protected by 2-factor authentication requires the correct password and the correct authentication of one of the registered trusted devices to verify the user's identity upon login. Note, that after the registration of a trusted device, the login might not be possible from clients where the device is not available, e.g. if a smartphone's integrated fingerprint reader is used as trusted device, the login might not be possible from other devices than the smartphone. In this case, a second trusted device, such as a USB key dongle, should be registered.

To complete the registration process, the user gets a verification email from our server to the submitted email address (our server uses *mSMTP* for sending emails). The email contains a one-time link that finally activates the user account. It is valid for 7 days, after which the account will automatically be deleted if it was not activated before. Anyway, the deletion is also possible manually before activation.

The email address is also required to reset a lost password. The user can click on the link "Forgot Password?" next to the password field and afterwards enter his email adress and a captcha code displayed as image, which is necessary to prevent abuse by bots. If the captcha code is correct, a new random password is sent to the user's email adress. The user now can sign in again and change his password.

The login process simply requires the registered email address and the password. If 2-factor authentication was activated for this account, access is authorised via one of the registered trusted devices afterwards. The user is now able to operate with the web service. Note, that a user is automatically logged out after 30 minutes of inactivity.

Users are able to change their email address and password or to delete their account at any time. In general, the deletion of an account implies the immediate removal of all data associated with this account. Specifically, the login credentials are removed from our database together with all uploaded files and data created for this user as a result from using the service.

#### 1.3 Job submission

A logged-in user may submit a new imputation job via our job submission form. Files can be provided for upload via three different ways: (a) directly in the browser, (b) via HTTP(s) URL or (c) via sFTP (see Sect. 1.3.1 for details). The user has to select the correct array build of his data (either GRCh37/hg19 or GRCh38/hg38) as this impacts the available reference panels for the correct build. We do not perform a lift over of the user's data, instead we provide all available reference panels in its lifted version as well, such that imputation output is always in the same build of the uploaded data. (See **Supplementary Table 1** for available reference panels.)

Further options are as described in the main paper and include the possibility to skip phasing or imputation, to select which information accompanies the haplotypes in the imputation output, an $R^2$-filter etc.

When submitting the job, the values in the form are translated to the command line options required for the *EagleImp* software. The relative location of the job folder, that is uniquely created for the input files, is submitted along with the command-line options to the job queuing system *Qmanager*.

The queuing status and progress of the job can be supervised in the *Jobs* section. Once a job has finished, the web service gets notified via a secured notification URL (allowing only connections from the backend system). According to the jobs return state (success or failed), the web service generated download links for the result files and notifies the user per email (see Sect. 1.4.1 for details).

##### 1.3.1 File upload

The web service offers three modes of file upload: Browser, URL and sFTP. During the upload files are checked on-the-fly to improve reliability and ensure stable job runs. In the case of a browser upload, a JavaScript module executed on the client's device checks selected files even before upload. Users get an immediate feedback about files not matching the required restrictions:

- A file must be a valid `*.vcf.gz` or `*.bcf` file.
- Only one file per chromosome is allowed, and each file must not contain the data of more than one chromosome.
- The file name must start with the chromosome identifier (in most cases a number, a leading "`chr`" is allowed).
- Allowed chromosome identifiers are numbers from 1 to 24, or "`X`" and "`Y`", but neither chromosome X is allowed to be submitted alongside chromosome 23, nor chromosome Y alongside 24.
- The *Pseudoautosomal Regions (PAR)* build an exception. Instead of a single file for chromosome 23/X a user may upload the PAR-regions separately. The identifiers have to be `23_PAR1`, `23_nonPAR` and `23_PAR2` then (whereby a leading `chr` is still allowed and `23` may be exchanged with `X`). (Note, that the service performs an automated split and merge if a single chromosome X file is uploaded.)
- The maximum file size must not exceed 1 GB (per file).
- The maximum number of samples must not exceed 100,000.

##### 1.3.2 Job queuing system

We implemented a simple and freely available job queuing software in the *Rust* programming language named *Qmanager* [5]. Its basic features are the reception of commands that should be processed in order on the backend system. It handles all queued, running and finished jobs (including their output and return codes) from the frontend system.

Job queuing request are handed over via a *https* connection in *JSON* format. For security reasons, the *Qmanager* is not allowed to run any arbitrary command. Accepted commands have to be registered in the configuration file of the *Qmanager* and are accessed via an identifier, not the command itself. Parameters though are passed through directly.

For *EagleImp-Web* the only registered command is the call to the customised *EagleImp* launch script. The script in turn launches the *EagleImp* executable for each of the user's input files (using a multi-processing scheme that launches several processes at once). Besides, the script also handles indexing of input and output files as well as PAR-region splitting of chromosome X. The parameters are the (relative) location of the user's input data (that is converted to an absolute path by the script) and the user options from the webinterface.

The *Qmanager* is run as a system service and provides options to submit a job to the queue, to remove jobs from the queue (either enqueued or finished) and to kill a running job. Whenever a job terminates (finished or killed), the *Qmanager* calls a pre-configured notification URL with the job identifier. In this case, the notification script is located on the frontend system and causes several operations such as changing the job state, notifying the user and preparing the download URLs.

A synchronised copy of the current job queue is permanently stored on the system's file storage. In the case of a system failure, the queue is automatically restored from that file. It is also possible to manually stop the queue without terminating it, e.g. for server administration. In this case, the currently running job will be regularly processed, but further jobs are delayed until the queue is put into the running state again. It is also possible to schedule new jobs.

#### 1.4 Job management

In the "Jobs" section users may supervise all recent job activities. For queued jobs the queuing state (current position in the queue) and submission time is displayed. Running jobs provide the current live progress detailed for each input file and the elapsed runtime. For finished jobs, the total runtime is shown and the result files are made available for download (see Sect. 1.4.1 for details).

Obviously, users may only see the status of their own jobs. The web service ensures that the reply of the *Qmanager* from a queue status request is filtered for the requesting user accordingly.

The current progress for each input file is read from a status file which is created by the *EagleImp* launch script. The *EagleImp* software continuously writes the progress of reading the input file and reference panel, phasing and imputation in this file, which is displayed by the web service as a progress bar. Additional information for each file is written by the *EagleImp* software in an info-, warning- and error-file, which contents are displayed by selecting the corresponding icon next to the progress bar.

The user may cancel a running job and delete queued, canceled or finished jobs immediately. By deleting a job, all input and output files from this job are removed from our server. Status and log files are still available to the user in the job history. By deleting a job from the history, all remaining data associated with this job is removed from our server. In order to save disk space on our server, input and output files of jobs will be deleted automatically after 7 days from completion, and the job is moved to the history. Jobs in the job history are removed automatically after a one year period if the user does not remove them manually. The automatic deletion is simply accomplished by a subprocess that is run on our server once a day. It deletes all input and output files of finished jobs that are older than 7 days and all other files and folders that are older than one year.

##### 1.4.1 Result download

A successful job run produces for each input file the following downloadable output files: the imputation output as `.imputed.vcf.gz`, the phasing output as `.phased.vcf.gz`, phasing confidences `.phased.confidences`, variant information `.varinfo` and the *EagleImp* log file `.log` (whereby some of these files may be optional depending on the user's selected job options). We also add *MD5* checksums (`.md5`) to the `.vcf.gz` files. We do not compress the imputation results again (e.g. as a `.zip`-file as for other imputation services) as `.vcf.gz` is already a compressed file format. The web service registers each downloadable file in the database and connects it with a unique random string. This string is formed to a unique download URL for this file and provided to the user as a download link. We use the *rewrite engine* by the Apache server to decode this URL to redirect a download request to our download engine, and the download engine queries the database for the associated file.

As we want to ensure that only authorised users are able to download their files, all files are locked per default, such that a user needs an active login prior to download a file (either directly in the browser or from the command line via *wget* or *curl*). Downloads from the command line can also be authorised via a unique password that is automatically generated for each finished job. The user receives this password together with the notification email for her/his job. This is advantageous if the imputation results have to be analyzed or post-processed on a different system than the user's workstation. The password has to be sent as a *POST* command after establishing the encrypted *https* connection to the web server to download a file. In detail, this is achieved by applying the `--post-data` option to send the password to the *wget* command. For convenience, we provide an automaticaly generated download script for each job (that uses *bash* and *wget*) to download all available files at once via the command line and which takes care about the authorization with the provided password.

### 2 Basic Concept of the *EagleImp* Software

This chapter provides a brief summary of the basic concept of the *EagleImp* software underlying our *EagleImp-Web* web service. For details, we refer to Wienbrandt *et al.* [3].

#### 2.1 Haplotype Phasing

The haplotypes of unphased genotypes from an input VCF-file are phased with *EagleImp* prior to imputation. Briefly summarised, for all genotypes, the most probable paths of all possible haplotype paths that explain the current target genotype are taken into account for determining the phase information of all heterozygous genotype calls. The detection of these paths is carried out by performing a *beam search.* The probability of a beam path which can be explained by the current genotype is the following:

$$P\left(h_{1\ldots m}^{\mathrm{mat}}, h_{1\ldots m}^{\mathrm{pat}} | g_{1\ldots m}\right) \approx \underbrace{P\left(g_{1\ldots m} | h_{1\ldots m}^{\mathrm{mat}}, h_{1\ldots m}^{\mathrm{pat}}\right)}_{\varepsilon^{n_{\mathrm{err}}}} P\left(h_{1\ldots m}^{\mathrm{mat}}\right) P\left(h_{1\ldots m}^{\mathrm{pat}}\right) \tag{1}$$

$h_{1\ldots m}$ is a haplotype sequence from positions 1 to $m$, $g_{1\ldots m}$ the respective target genotype sequence. $n_{\mathrm{err}}$ is the number of consistency errors between the pair of haplotype paths to the target genotypes and $\varepsilon$ is a fixed error probability. $P\left(h_{1\ldots m}^{\mathrm{mat}}\right)$ and $P\left(h_{1\ldots m}^{\mathrm{pat}}\right)$ are the maternal and paternal haplotype path probabilities which are calculated as follows:

$$P\left(h_{1\ldots m}\right) \approx \sum_{x=m-H}^{m-1} P\left(h_{1\ldots x}\right) f\left(h_{x+1\ldots m}\right) P\left(\mathrm{rec}\ m|x\right) \tag{2}$$

This equation includes the calculation of the (relative) frequency $f$ of a sequence $h_{x+1\ldots m}$ in the reference. ($H$ is a history parameter and $P\left(\mathrm{rec}\ m|x\right)$ is the recombination probability between two sites $m$ and $x$). The probability to keep a phase at a certain heterozygous position in the target is simply calculated by the relation of all beam paths at this position that also keep the phase to all beam paths with any phase. So far, this is the same procedure as in *Eagle2* [7].

*EagleImp* implements a simple forward phasing step for determining the initial phases and phasing confidences, and performs a subsequent reverse phasing step to refine the decisions from the forward step.

Phasing preliminaries include the creation of a *condensed reference* for each target that includes (a) only the best $K$ haplotypes from the reference compared to the current target ($K$ is a runtime parameter), (b) the haplotype information from the reference of those $K$ haplotypes only at heterozygous positions, and (c) the information for segments between heterozygous positions if the segment in the haplotype is consistent with the current target or not. The condensed reference is the basis for a *PBWT* data structure that enables fast lookups of the frequency $f$ from Eq. 2 (with a constant runtime complexity).

As targets are independent of each other, the phasing process is trivially parallelised by processing several targets concurrently distributed over the available system threads.

#### 2.2 Genotype Imputation

As in the phasing step the main data structure used in the imputation algorithm is the *PBWT*. Note, that this equals the name of the original imputation tool *PBWT* [8] where this data structure was introduced first and which forms the base for the imputation part here.

The first step in *PBWT* imputation is to determine the *set-maximal matches* from a target haplotype sequence to the complete reference. A match is set-maximal if there is no longer match including this one, and there is also no other sequence in the reference with a longer match that includes this region. The matches are used to calculate the imputation allele dosage for the variants in the gaps between call sites that need to be imputed. This is done by a simple scoring function that uses all matches that cover a gap and weighs the length of a match (the longer the better) and the position of the match in the gap (the further from the edges the higher the score). Let $m$ be the position of the last call site before the gap and $m_i^{\text{start}}$ and $m_i^{\text{end}}$ the positions of an overlapping match with index $i$. The score of match $i$ is then calculated as follows.

$$s_i = \left(m - m_i^{\text{start}} + 1\right)\left(m_i^{\text{end}} - m\right) \tag{3}$$

The allele dosages of the variants in the gap at $m$ are simply calculated as the scores at $m$ where the corresponding allele in the reference is the alternative allele in relation to all scores at $m$. From the allele dosages other variables such as genotype dosages, genotype probabilities, imputation $r^2$ etc. are calculated.

As in the phasing step, targets are also independent in the imputation step. The calculation of set-maximal matches is therefore done separately for each target whereby the targets are distributed over the available system threads. However, the VCF output format is in variant-major format, which implies that all targets have to be imputed at least up to the current output line before the output can be written. Furthermore, this process is dominated by high IO requirements but low CPU load. So, a typical parallelisation over the targets and imputing the targets at once is not practical. In *EagleImp* [3] we developed a multi-processing scheme that uses several threads to impute only chunks from each target in parallel and writes to several temporary output files concurrently. This way, we were able to widen the IO bottleneck and can efficiently take advantage of the multi-processing capabilities of our computing system.

### 3 Integration of FPGA Acceleration in *EagleImp*

The *EagleImp* phasing preliminaries including the creation of the PBWT data structure require a large amount of computational resources but mainly consist of operations based on Boolean values, that are highly suitable for the implementation on a *field-programmable gate array (FPGA)*. We implemented an FPGA design to outsource this part to our Alpha Data ADM-PCIE-8K5 FPGA accelerator card (equipped with a Xilinx Kintex UltraScale KU115 FPGA, two attached 8 GB SODIMM memory modules and a PCI Express Gen3 x8 high-speed connection to the host). We implemented 32 parallel pipelines for the concurrent processing of 32 targets in our FPGA design. We divided a pipeline into two major parts: (1) creation of the condensed reference with (2) subsequent generation of the PBWT structure, whereby a certain pre- and post-processing is required on the host system as well. For preparing the source code (written in the *VHDL* hardware description language) we used *Sigasi Studio XL* [9], and we compiled the FPGA configuration bitfile with *Xilinx Vivado 2020.2* [10]. An overview of the processing pipeline is shown in **Figure 2** in our main paper. We explain the illustrated steps in the following.

#### 3.1 Target Preparation

As the FPGA design contains 32 parallel pipelines for concurrent target processing, the host system divides all targets in blocks of 32 target samples for transmission to the FPGA. As in CPU-only phasing, the host determines the call sites and the $K$-best reference haplotypes for each target. The FPGA is then initialised with a number of constants (mainly the size of the targets and the reference panel). The data for each target provided by the host contains the complete genotype information of the targets (encoded as two bitstreams with the first indicating the genotypes that are the homozygous reference type ("is 0"), and the second indicating the genotypes that are homozygous alternative type ("is 2")), the call sites (encoded as a bitstream with one bit for each site where a set bit indicates a call site), and a bit mask for the $K$ best haplotypes (i.e. one bit for each haplotype with a set bit indicating that this haplotype is to be used in the analysis). The FPGA distributes the target data from the host to the FPGA pipelines in a simple round-robin scheme, whereby a block of targets has to be processed completely before the next block can be initialised.

#### 3.2 Creation of the Condensed Reference

After the initialization of all pipelines with the target data, the host streams the reference data to the FPGA in variant-major format, i.e. for each variant the haplotype data of all samples is provided. The reference data has to be streamed completely repeated for each block of targets. The FPGA processes one variant after the other. The reference data is broadcast to all pipelines in parallel with 16 bits of haplotype information per clock cycle. The *checkIncon* unit produces an *inconsistency stream* by checking the consistency of the 16 presented haplotypes to the current genotype according to the following equation:

$$\texttt{inc} = (\texttt{is0}\ \mathrm{AND}\ \texttt{hap})\ \mathrm{OR}\ (\texttt{is2}\ \mathrm{AND}\ \mathrm{NOT}\ \texttt{hap}) \qquad (4)$$

An execption is made for variants that are marked as a call site. The haplotype data of these variants is forwarded unchanged.

The subsequent unit selects the *K-best haplotypes* from the incoming stream. For each incoming 16bit-word the corresponding 16bit-mask is used to remove the unneeded haplotypes. As for the follow-up processes it is required to reduce the stream of $n$ bits for each variant (one bit for each reference haplotype) to only $K$ bits, the unit reduces the incoming 16bit-words to words of a size that matches the number of set bits of the current mask. This is achieved by using a routing similar to an *odd-even transposition sort* sorting network, where the mask is sorted and the data routed alongside the 1-bits of the mask. The resulting smaller words are then collected until full 16bit-words are available, which are provided to the *condense* unit afterwards.

The *condense* unit collects the information for all variants that are not marked as call sites (which is the current inconsistency information) in a local BRAM that is able to store $K$ bits. The data for all subsequent variants that are not call sites are merged with the currently stored data by a simple logical OR operation in order to generate the information for the current inconsistency segment in the condensed reference (refer to Wienbrandt *et al.* [3] for more details on the condensed reference). This process continues until a variant is marked as a call site which triggers the release of the data stored in the local BRAM (which is the complete inconsistency information for the current segment) and the incoming data from the call site to be stored. The next variant triggers the release of the just stored data for the call site while the incoming data is now stored in the BRAM.

The outgoing condensed reference is then buffered before being converted to a PBWT structure in the next step. The buffer is able to block the preliminary processes to ensure no data loss in the case the buffer runs full. Furthermore, it ensures the correct crossing of clock domains as the creation of the condensed reference runs with a different clock frequency than the following PBWT creation.

#### 3.3 Creation of the PBWT

The required PBWT structure for the phasing process differs from the so far created condensed reference only in the permuted haplotype information at each site (such that the original haplotypes appear sorted when read in the permuted order backwards starting from that site). For details of the PBWT and the original creation algorithm, we refer to Wienbrandt *et al.* [3] and Durbin [8].

To generate the permuted haplotypes the PBWT unit contains two haplotype buffers for storing up to $K$ bit of haplotype data each, and two permutation arrays that can store up to $K$ permutation indices (of size $\log_2 K$). The buffers require access to an arbitrary stored haplotype bit preferably in a single clock cycle, which is implemented in the FPGA's local BRAM. (In fact, the data is available in the next clock cycle after the required address has been provided, but the process is pipelined such that each clock cycle a different address can be provided with the output appearing with one cycle delay.)

The PBWT generation process is performed as follows. First, the incoming condensed reference data (in 16bit words) is serialised to a bitwise stream. The data for the first site is simply stored in the first haplotype buffer at the beginning, and the first permutation array is initialised with the identity function (i.e. no permutation will be made). Meanwhile, for each site, the number of zero bits (*count0*) is determined from the serialised stream. For the following sites the process is as follows. The bitstream is stored in the second haplotype buffer while the first buffer is read according to the permutation stored in the first permutation array. In detail, the permutation indices are read subsequently and provided as address to the haplotype buffer. The resulting data stream is now in the required permuted order for the compact PBWT representation and is provided to a deserialisation unit that prepares 32bit output words.

To generate the next permutation, the second permutation array is implemented as two queues: the first one starting at the beginning of the array for storing the permutation indices that addressed a zero bit in the haplotype buffer, and the second one starting at *count0* for storing the indices that addressed a one bit. Each current permutation index is then written to the second (new) permutation array with its destination dependent on the corresponding haplotype bit: if the bit is zero the index is written to the first queue, if it is one it is written to the second queue. After the last data from the current site all arrays are switched before the next site, i.e. the first haplotype buffer becomes the second and vice versa, and the first permutation array becomes the second and vice versa. The design presented here is the exact hardware implementation of the algorithm listing for compact PBWT creation provided in Wienbrandt *et al.* [3].

The generation of the compact PBWT is the major bottleneck of our FPGA design as the complete condensed reference needs to be processed bit by bit. We were able to increase the data throughput by processing two bits of the condensed reference in one clock cycle. For this, we used the dual-port feature of the FPGA's BRAM units. In detail, two permutation indices are read from the first permutation array in each clock cycle. Both indices request a data bit from the haplotype buffer from different addresses in the same clock cycle, which is only possible with the dual-port feature. The resulting two data bits generate four possibilities of writing the two indices to the second permutation array: (a) both indices get into the first queue, (b) the first index gets into the first queue and the second in the second queue, (c) the first gets into the second queue and the second gets into the first queue, and (d) both indices get into the second queue. Again, this is only possible using the BRAM's dual-port feature.

For a maximum throughput the design's clock frequency is a major factor. We are able to operate the PBWT creation at an increased frequency of 266 MHz while the rest of the pipeline operates at 114 MHz. This is a factor of 2.33 times faster but while the PBWT unit processes only 2 bits per clock cycle and the condensed reference is provided with 16 bits per clock cycle, we can predict the throughput of the PBWT unit to be $\frac{16}{2} \times \frac{114}{266} = 3.43$ times slower than the maximum throughput of the condensed reference unit. However, because the condensed reference unit provides its output at a different (in general lower) speed than the input of the raw data is processed (as only call sites generate sites in the condensed reference and the reference is reduced to the $K$ best haplotypes only), the bottleneck is reduced or even eliminated in most cases making the FPGA work at a maximum efficiency.

#### 3.4 Data Collection and Post Processing

Before sending the PBWT data to the host system it is buffered in the attached DRAM. The output words (32 bit) from each deserialisation unit are collected to form 512 bit words according to the size of the DRAM data port. Due to the processing width of 2 bits at a frequency of 266 MHz the PBWT unit can generate a 512bit word only every 256 clock cycles, i.e. only every 110 clock cycles in the 114 MHz clock domain. This leaves enough space for the DRAM to handle the output of the 32 parallel pipelines without generating another bottleneck. We store the compact PBWT data of each target ordered in subsequent memory areas in the DRAM, each area designated to one pipeline. The size of the memory area depends on the number of expected DRAM words for the designated target PBWT and is simply achieved by adding an address offset to each pipeline corresponding to the number of expected DRAM words for all preliminary pipelines.

As our *Alpha Data* FPGA accelerator board is equipped with two DRAM modules, we implemented two DRAM buffers as well. While one buffer is filled by the processing pipelines for the current target block, the other buffer is read and the data from the previous target block is transferred to the host system simultaneously.

The host system generates the index fields required for a fast access to the PBWT data on-the-fly while copying the received data from the transfer buffer to host memory. The index is simply implemented as a 32bit integer attached to each 32bit block of compact PBWT data. The integer value is the number of zero bits from the beginning of the current site up to the current block (inclusive) (see Wienbrandt *et al.* [3] for details on the use of the index). We decided against the implementation on the FPGA as this doubles the amount of data to be transferred to the host which we do not want to risk to become the bottleneck of the design. Furthermore, this process is very efficiently implemented on the host by using the *popcount* processor directive that determines the number of set bits in a 32bit word in a single processor command.

##### 3.5 Design Limitations

From the FPGA's hardware resources the local BRAM is the main limiting factor. The most BRAM resources are required to store the intermediate permutation arrays (current and new one) during PBWT creation. Their size is directly related to the size of the parameter $K$ as $K$ indices need to be stored in each array. We therefore decided to limit the maximum value of $K$ to 32,768 in our design to be able to implement 32 processing pipelines on the FPGA. We are planning to reorganise BRAM resources to be able to implement a higher possible value of $K$ in the future, e.g. by reducing the number of pipelines to 16 the maximum value of $K$ could be increased to 65,536 easily, but we expect a reduction in the processing speed of the FPGA in the same manner, which, however, is subject to further analyses.

#### 4 Supplementary Tables and Figures

Supplementary Table 1: The reference panels available in *EagleImp-Web* are based on the *1000 Genomes Phase 3* release [11] and the EGA release of the *HRC1.1* panel [12, 13].

| Build | Reference | #Variants | #Samples | Notes |
|---|---|---|---|---|
| GRCh37/hg19 | 1000G Phase 3 | 84,801,880 | 2,504 | native release |
| | HRC1.1 EGA release | 40,405,505 | 27,165 | native release |
| GRCh38/hg38 | 1000G Phase 3 | 82,499,492 | 2,504 | native release |
| | 1000G Phase 3 lifted | 84,707,528 | 2,504 | liftover hg19 to hg38 |
| | HRC1.1 EGA release lifted | 40,388,400 | 27,165 | liftover hg19 to hg38 |

Supplementary Table 2: Benchmark datasets for quality assurance and runtime measures with FPGA acceleration in *EagleImp-Web*. The datasets are the same as used in Wienbrandt *et al.* [3] for benchmarking the *EagleImp* software.

| | Target | #Variants | #Samples | Ancestry |
|---|---|---|---|---|
| (1) | *HRC.EUR* | 619,872 | 494 | European |
| (2) | *COVID.Italy* | 559,519 | 2,113 | Italian |
| (3) | *COVID.Spain* | 549,696 | 1,792 | Spanish |

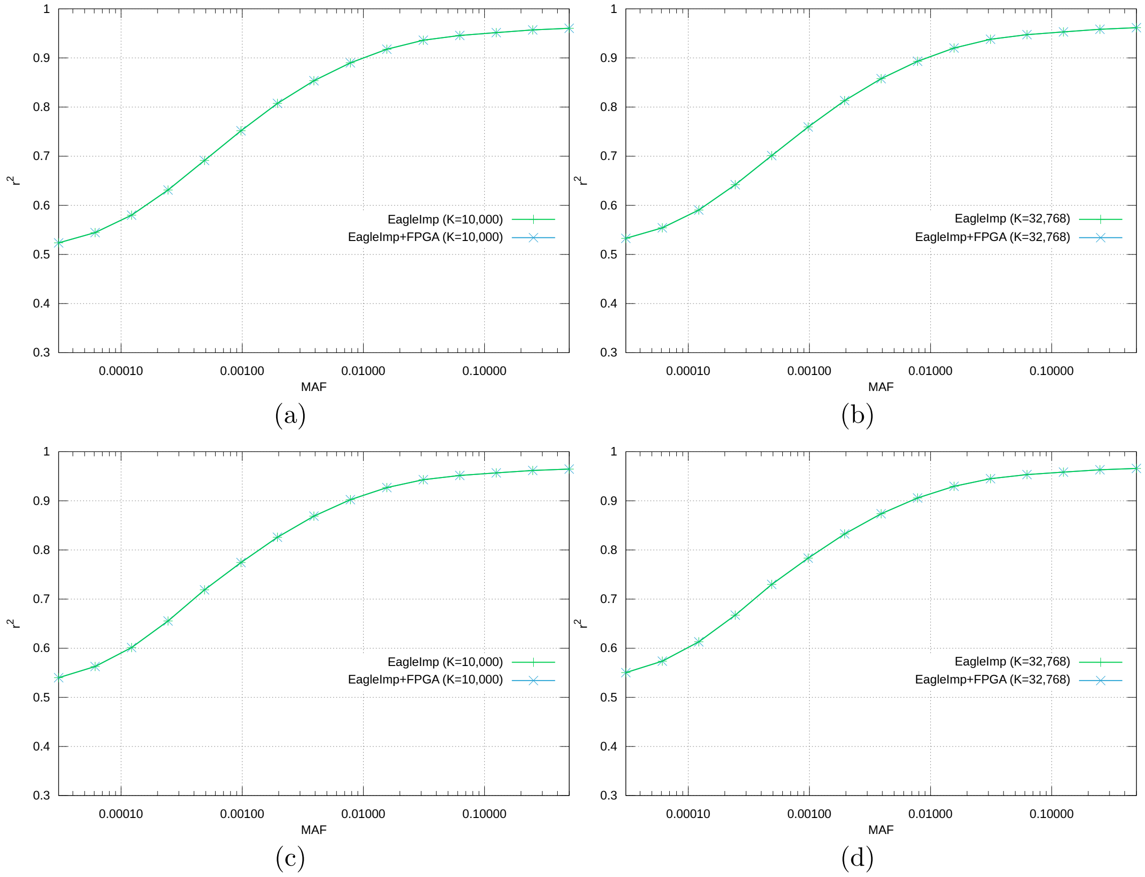

Supplementary Figure 1: Imputation accuracy $r^2$ stratified by minor allele frequency (MAF) for $K = 10,000$ and $K = 32,768$ for two real-world datasets *COVID.Italy* (a–b) and *COVID.Spain* (c–d) imputed using the EGA release of the HRC1.1 reference panel. The runs with and without FPGA acceleration do not show any difference demonstrating that there is no impact on imputation quality when using FPGA acceleration for the preliminary phasing step in *EagleImp*.

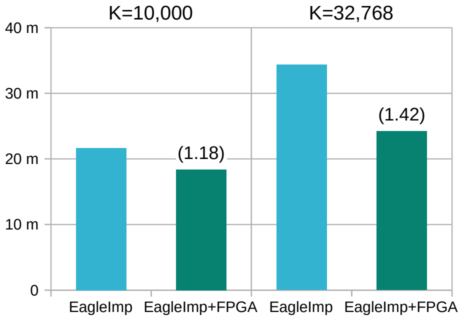

Supplementary Figure 2: Wall-clock runtimes in seconds of benchmarks with the *HRC.EUR* dataset comprising 494 European samples extracted from the EGA release of the HRC1.1 reference panel, reduced to 647,963 variants (from Illumina's *Global Screening Array (GSA)* and imputed against the remaining samples from the same panel with two different $K$ parameters. Numbers in brackets indicate the speedup of the FPGA-accelerated run when compared to the CPU-only run of *EagleImp*.

Supplementary Table 3: Wall-clock runtimes in seconds of benchmarks with the *HRC.EUR* dataset comprising 494 European samples extracted from the EGA release of the HRC1.1 reference panel, reduced to 647,963 variants (from Illumina's *Global Screening Array (GSA)* and imputed against the remaining samples from the same panel with two different $K$ parameters.

| *HRC.EUR* | *EagleImp* (CPU-only) | *EagleImp* (FPGA-accelerated) | Speedup |
|---|---|---|---|
| $K$=10,000 | 1299 | 1104 | 1.18 |
| $K$=32,768 | 2061 | 1453 | 1.42 |

Supplementary Table 4: Wall-clock runtimes in seconds for the GWAS benchmark dataset *COVID.Italy* consisting of 2,113 Italian samples with 559,519 variants imputed with *EagleImp* against the HRC1.1 EGA release, using two different values of the phasing parameter $K$.

| *COVID.Italy* | *EagleImp* (CPU-only) | *EagleImp* (FPGA-accelerated) | Speedup |
|---|---|---|---|
| $K$=10,000 | 5240 | 4257 | 1.23 |
| $K$=32,768 | 8620 | 5272 | 1.64 |

Supplementary Table 5: Wall-clock runtimes in seconds for the GWAS benchmark datasets *COVID.Spain* consisting of 1,792 Spanish samples with 549,696 variants imputed with *EagleImp* against the HRC1.1 EGA release, using two different values of the phasing parameter $K$.

| *COVID.Spain* | *EagleImp* (CPU-only) | *EagleImp* (FPGA-accelerated) | Speedup |
|---|---|---|---|
| $K$=10,000 | 4324 | 3473 | 1.25 |
| $K$=32,768 | 7083 | 4278 | 1.66 |

### 5 Exemplary COVID-19 GWAS Imputed with *EagleImp-Web*

For an exemplary GWAS we replicated our findings of our COVID-19 association study from Ellinghaus, 2020 [14]. The corresponding datasets *COVID.Italy* and *COVID.Spain* comprise 2113 Italian samples (839 cases, 1,274 controls) typed at 559,519 variants and 1792 Spanish samples (842 cases, 950 controls) typed at 549,696 variants respectively. We performed imputation of these datasets using the EGA release of the HRC1.1 reference panel [12, 13] and three different $K$ parameters: $K = 10,000$, $K = 32,768$ and $K = \max$. Subsequently, we applied our *BIGwas* association pipeline [15] on the imputed results. The resulting Manhattan plots are depicted in the following **Supplementary Figures 3 (a–c)**.

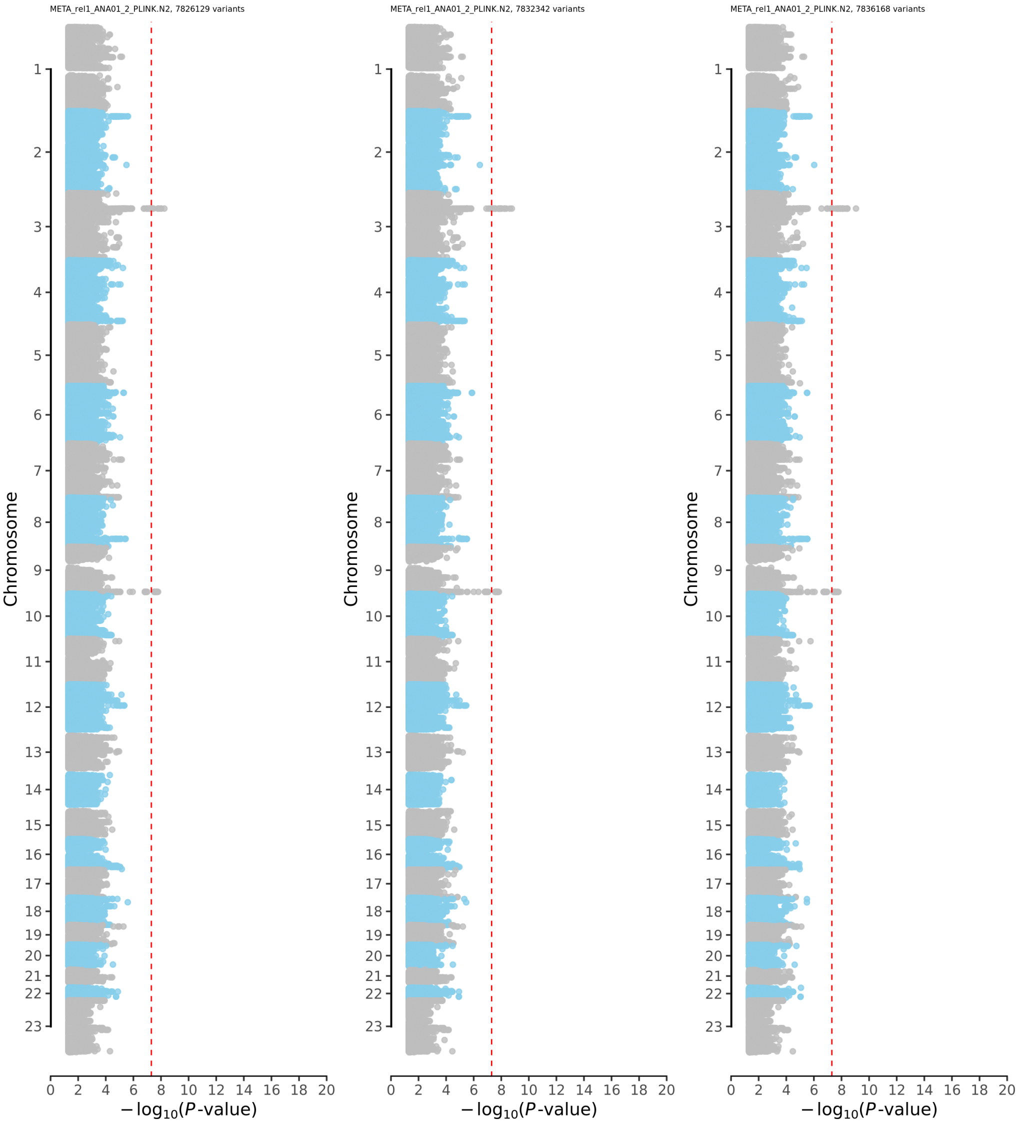

(a) (b) (c)

Supplementary Figure 3: COVID-19 GWAS summary (Manhattan) plots of three meta-analyses of *COVID.Italy* and *COVID.Spain* association statistics after phasing and imputation with *EagleImp-Web* and three different $K$ parameters: (a) $K = 10,000$, (b) $K = 32,768$ and (c) $K = \text{max}$. We used exactly the same GWAS input datasets *COVID.Italy* and *COVID.Spain* for phasing and imputation as in [14]. The dashed red line shows the genome-wide significance threshold at $P = 5 \times 10^{-8}$. The variants were filtered with an imputation accuracy $r^2$-limit of 0.6 (as done in [14]). The figures show the replication of genome-wide significant association findings from our COVID-19 GWAS [14] and indicate smaller (i.e. more significant) genome-wide significant P-values of GWAS lead variants at the established COVID-19 risk locus 3p21.31 with higher values of $K$ ($P = 5.72 \times 10^{-9}$ for $K = 10,000$, $P = 1.71 \times 10^{-9}$ for $K = 32,768$, $P = 9.91 \times 10^{-10}$ for $K = \text{max}$)

#### References

[1] Apache HTTP Server Project, `https://httpd.apache.org`.

[2] PostgreSQL: The World's Most Advanced Open Source Relational Database, `https://www.postgresql.org`.

[3] Wienbrandt, L. and Ellinghaus, D. (2022) EagleImp: Fast and Accurate Genome-wide Phasing and Imputation in a Single Tool. *bioRxiv,*.

[4] Wienbrandt, L. EagleImp, `https://github.com/ikmb/eagleimp.git`.

[5] Kässens, J. C. Qmanager, `https://github.com/ikmb/qmanager.git`.

[6] A simple PHP WebAuthn (FIDO2) server library, `https://github.com/lbuchs/WebAuthn`.

[7] Loh, P.-R., Danecek, P., Palamara, P. F., Fuchsberger, C., Reshef, Y. A., Finucane, H. K., Schönherr, S., Forer, L., McCarthy, S., Abecasis, G. R., et al. (October, 2016) Reference-based phasing using the Haplotype Reference Consortium panel. *Nature Genetics,* **48**, 1443–1448.

[8] Durbin, R. (2014) Efficient haplotype matching and storage using the positional Burrows-Wheeler transform (PBWT). *Bioinformatics,* **30**(9), 1266–1272.

[9] Sigasi;, `https://www.sigasi.com`.

[10] Xilinx Vivado, `https://www.xilinx.com/products/design-tools/vivado.html`.

[11] The 1000 Genomes Project Consortium (September, 2015) A global reference for human genetic variation. *Nature,* **526**, 68–74.

[12] McCarthy, S., Das, S., Kretzschmar, W., Delaneau, O., Wood, A. R., Teumer, A., Kang, H. M., Fuchsberger, C., Danecek, P., Sharp, K., et al. (August, 2016) A reference panel of 64,976 haplotypes for genotype imputation. *Nature Genetics,* **48**(10), 1279–1283.

[13] European Genome-Phenome Archive Haplotype Reference Consortium Release 1.1 - subset for release via the EGA, `https://ega-archive.org/datasets/EGAD00001002729`.

[14] Ellinghaus, D., Degenhardt, F., Bujanda, L., Buti, M., Albillos, A., Invernizzi, P., Fernández, J., Prati, D., Baselli, G., Asselta, R., et al. (October, 2020) Genomewide association study of severe COVID-19 with respiratory failure. *The New England Journal of Medicine,* **383**(16), 1522–1534.

[15] Kässens, J. C., Wienbrandt, L., and Ellinghaus, D. (June, 2021) BIGwas: Single-command quality control and association testing for multi-cohort and biobank-scale GWAS/PheWAS data. *Gigascience,* **10**(6), giab047.
